# Quantitative Trait Loci Controlling *Phytophthora cactorum* Resistance in the Cultivated Octoploid Strawberry (*Fragaria* x *ananassa*)

**DOI:** 10.1101/249573

**Authors:** Charlotte F. Nellist, Robert J. Vickerstaff, Maria K. Sobczyk, César Marina-Montes, Philip Brain, Fiona M. Wilson, David W. Simpson, Adam B. Whitehouse, Richard J. Harrison

## Abstract

The cultivated strawberry, *Fragaria x ananassa* (*Fragaria* spp.) is the most economically important global soft fruit. *Phytophthora cactorum*, a water-borne oomycete causes economic losses in strawberry production globally. A bi-parental cross of octoploid cultivated strawberry segregating for resistance to *P. cactorum*, the causative agent of crown rot disease, was screened using artificial inoculation. Multiple resistance quantitative trait loci (QTL) were identified and mapped. Three major effect QTL (*FaRPc6C*, *FaRPc6D* and *FaRPc7D*) explained 36% of the variation observed and in total, the detected QTL explained 86% of the variation observed. There were no epistatic interactions detected between the three major QTLs. Testing a subset of the mapping population progeny against a range of *P. cactorum* isolates revealed no major differences in host response, however, some lines showed higher susceptibility than predicted, indicating that additional undetected factors may affect the expression of some quantitative resistance loci. Using historic crown rot disease score data from strawberry accessions, a preliminary genome-wide association study of 114 individuals revealed additional loci associated with resistance to *P. cactorum*. Mining of *Fragaria vesca* Hawaii 4 v1.1 genome revealed candidate resistance genes in the QTL regions.

## INTRODUCTION

The cultivated strawberry, *Fragaria x ananassa* (*Fragaria* spp.) is the most economically important global soft fruit and is an integral part of the diet of millions of people ^1^. In a recent modelling study, a reduction in fruit and vegetable production or a reduction in affordability due to climate change has been predicted to be a key driver of food insecurity. Twice as many climate-related deaths were associated with reductions in fruit and vegetable consumption than with climate-related increases in the prevalence of underweight individuals ^2^.

Traditionally, the major strategy for disease control in strawberry production relied heavily upon pre-plant fumigation and chemicals. The withdrawal of methyl bromide along with other active chemicals, including fungicides and soil fumigants are increasing the challenges in field strawberry production, resulting in a rise of occurrences and severities of some once well controlled diseases ^3^. A switch to producing strawberries in soilless substrate is now common practice across the world. The soilless substrate system offers many advantages, including the benefit of separating the strawberries from the infected soil ^4^. This has resulted in a reduction in the prevalence of some soil-borne diseases, but not for water-borne pathogens such as the hemibiotrophic oomycete, *Phytophthora cactorum. P. cactorum* (Lebert and Cohn) Schröeter is a destructive pathogen, that can infect a wide variety of plant species, causing serious damage in both ornamental and agricultural crops ^5^. It is the causative agent of strawberry crown rot ^6^ and strawberry leather rot ^7^, affecting the fruit. Both diseases are reported to cause economic losses in strawberry production globally; in Norway in 1996/97 there were reports of plants losses of up to 40% caused by crown rot ^8^ and in 1981 reports of commercial farms in Ohio described crop losses from leather rot of 20-30% ^9^. Amplified fragment length polymorphism (AFLP) analysis of *P. cactorum* isolates of crown rot and leather rot showed they are distinctly different from each other and from *P. cactorum* isolated from other hosts ^10^. No correlation has been found between resistance to crown rot and resistance to leather rot ^11^.

Strawberry plants infected with *Phytophthora* crown rot develop initial symptoms in spring and summer, frequently during hot periods. Plants can often appear stunted; the youngest leaves are usually the first to wilt, followed by the older leaves, eventually resulting in the collapse and death of the plant ^12^. Red-brown lesions and longitudinal splits can be observed within the crown ^13^. Sexually produced oospores are the primary source of inoculum; these are the resting spores that can persist in the soil or infected plants for many years ^12^. Under the conducive conditions of saturated soil, oospores germinate to produce sporangia which release the motile asexual life stage, zoospores. Zoospores are chemotactically attracted to nearby roots ^14^, where they attach to the root surface, encyst and penetrate the root epidermis.

The public breeding programme at East Malling (NIAB EMR, Kent, UK), since its establishment in 1983, has successfully released 43 strawberry cultivars to the Northern European market. Efforts have focused on combining excellent fruit quality with high yield, low percentage waste and resistance to filamentous diseases. Breeding for disease resistance is a high priority for many breeding programmes across the world. There has been extensive research investigating qualitative (major gene) resistance to *Phytophthora* species (for a selection of *R* gene - *Avr* gene interactions see Table 2 in Vleeshouwers *et al.* ^15^), however, much less is known about quantitative resistance (multiple genes, each of partial effect) to *Phytophthora* species. Quantitative trait loci (QTL) mapping is a routine technique for pinpointing genes controlling complex polygenic traits to specific regions of the genome, through statistical analysis. Previous studies have identified resistance to *P. cactorum* in the octoploid strawberry and it appears to be under polygenic control ^16–18^, with a major locus, *FaRPc2*, recently reported on linkage group 7D ^19^. Variation in resistance has been also observed in the wild progenitors of *F*. x *ananassa*; *Fragaria chiloensis* and *Fragaria virginiana* populations ^20^.

**Table 2.** Evaluation of 15 representative individuals of the ‘Emily’ x ‘Fenella’ population and their resistance/susceptibility response to *Phytophthora cactorum* isolates, with presence/absence of resistance quantitative trait loci (QTL) detailed.

| Individual | Predicted Score | Phytophthora cactorum isolate |  |  |  | Presence/absence of resistance quantitative trait loci (QTL) |  |  |  |  |  |  |  |  |  |  |  | Marker Total |
| --- | --- | --- | --- | --- | --- | --- | --- | --- | --- | --- | --- | --- | --- | --- | --- | --- | --- | --- |
|  |  | P414 | P404 | P415 | P416 | FaRPc1B | FaRPc2C | FaRPc3B | FaRPc3C-A | FaRPc3C-B | FaRPc5B | FaRPc6A | FaRPc6B | FaRPc6C | FaRPc6D | FaRPc7A | FaRPc7D |  |
| EF147 | 1.3 | 1.9 | 2.6 | 2.7 | 1.4 | 0 | 1 | 1 | 1 | 0 | 1 | 1 | 1 | 1 | 1 | 1 | 1 | 10 |
| EF011 | 1.6 | 1.0 | 2.3 | 1.6 | 2.3 | 1 | 1 | 1 | 1 | 0 | 0 | 1 | 1 | 1 | 1 | 0 | 1 | 9 |
| EF101 | 1.6 | 1.6 | 2.6 | 2.8 | 2.0 | 0 | 1 | 0 | 1 | 0 | 1 | 1 | 1 | 1 | 1 | 1 | 1 | 9 |
| EF184 | 2.2 | 1.7 | 3.0 | 3.6 | 1.7 | 0 | 1 | 1 | 1 | 0 | 0 | 0 | 0 | 1 | 1 | 1 | 1 | 7 |
| EF021 | 2.5 | 1.9 | 2.9 | 2.3 | 4.0 | 1 | 1 | 1 | 0 | 1 | 1 | 0 | 0 | 1 | 1 | 0 | 0 | 7 |
| EF141 | 2.9 | 3.4 | 2.0 | 4.0 | 4.3 | 1 | 0 | 1 | 1 | 1 | 0 | 0 | 1 | 0 | 0 | 1 | 1 | 7 |
| EF164 | 2.7 | 3.0 | 3.8 | 2.7 | 4.0 | 1 | 1 | 0 | 0 | 0 | 1 | 0 | 0 | 0 | 1 | 1 | 1 | 6 |
| EF041 | 3.4 | 4.0 | 3.6 | 3.4 | 3.7 | 0 | 0 | 1 | 0 | 1 | 1 | 0 | 0 | 0 | 0 | 1 | 1 | 5 |
| EF060 | 3.8 | 3.2 | 3.2 | 2.0 | 2.7 | 1 | 0 | 0 | 0 | 0 | 1 | 0 | 0 | 1 | 0 | 1 | 0 | 4 |
| EF187 | 3.7 | 3.1 | 2.7 | 2.7 | 2.3 | 0 | 0 | 0 | 1 | 0 | 0 | 1 | 1 | 0 | 0 | 0 | 1 | 4 |
| EF120 | 4.4 | 6.0 | 6.4 | 6.4 | 6.6 | 0 | 0 | 0 | 1 | 1 | 0 | 0 | 1 | 0 | 0 | 0 | 0 | 3 |
| EF166 | 4.2 | 5.9 | 6.6 | 6.5 | 6.6 | 0 | 0 | 0 | 0 | 0 | 1 | 0 | 1 | 1 | 0 | 0 | 0 | 3 |
| EF035 | 4.7 | 5.5 | 5.8 | 5.3 | 4.0 | 0 | 0 | 1 | 0 | 0 | 1 | 0 | 0 | 0 | 0 | 0 | 0 | 2 |
| EF084 | 4.7 | 5.6 | 6.3 | 5.0 | 4.0 | 0 | 0 | 0 | 0 | 0 | 1 | 0 | 1 | 0 | 0 | 0 | 0 | 2 |
| EF040 | 4.9 | 5.4 | 5.1 | 5.4 | 3.9 | 0 | 0 | 0 | 0 | 0 | 1 | 0 | 0 | 0 | 0 | 0 | 0 | 1 |

The cultivated strawberry (2*n* = 8*x* = 56) is an allo-polyploid outbreeder with a genome comprised of four comparable homeologous sets of diploid chromosomes ^21,22^. The octoploid genome is estimated to be 698 Mb; 80% of the size of quadrupling the diploid genomes (~200 Mb each) ^23^. The development of the 90K SNP (single nucleotide polymorphism) Affymetrix® IStraw90 Axiom® Array ^24^ has aided genetic studies and marker-assisted breeding. The genomes of 19 octoploid and six diploid strawberry accessions were sequenced to serve as resources for SNP discovery and interpretation. The octoploids used were ‘Holiday’, ‘Korona’, ‘Emily’, ‘Fenella’, ‘Sweet Charlie’, ‘Winter Dawn’, ‘CA65.65.601’ and ‘NH-SB480’, six F_1_ progeny of ‘Holiday’ x ‘Korona’, one F_2_ progeny of ‘Dover’ x ‘Camarosa’, one *F. virginiana* and three *F. chiloensis* ^24^. The diploids used were the known progenitor *Fragaria vesca* and two probable progenitors *Fragaria iinumae* and *Fragaria mandshurica* ^24^. A high percentage of SNP markers were designed sub-genome-specific to detangle the complexity of the allo-octoploid genome and allow accurate scoring. However, its widespread use was limited by cost. A smaller, cheaper version of the array, Axiom® IStraw35 384HT, has been developed by combining mapped SNP probes from multiple groups from across the world and contains just over 34 000 markers ^25,26^.

The genus *Phytophthora* comprises of numerous destructive crop pathogens ^27^. The most extensively studied are *Phytophthora infestans* (late blight of potato and tomato) and *Phytophthora sojae* (root and stem rot of soybean). The majority of *R* genes identified in potato against *P. infestans* belong to the coiled-coil, nucleotide-binding, leucine-rich repeat (CC-NLR) class of intracellular immune receptors ^28^. The corresponding avirulence (*Avr*) genes identified belong to the RxLR class of effectors ^28^. These secreted, modular effectors have an RxLR motif for translocation into the host cell with a quickly evolving effector domain at the C-terminus. Fewer extracellular resistance genes have been characterised, however, the few that have, have been associated with resistance to multiple *Phytophthora* pathogens. Cell surface L-type-lectin-RLKs (receptor-like kinases) have been associated with resistance to *Phytophthora brassicae*, *Phytophthora capsici* and *P. infestans*^29-31^. An extracellular receptor-like protein (RLP) ELR (elicitin response) was identified in a wild species of potato. ELR was found to recognise elicitin proteins from a diverse set of *Phytophthora* species, including *P. infestans*, *P. sojae* and *Phytophthora cryptogea* ^32,33^.

In this study, the genetic basis of quantitative resistance to *P. cactorum* was investigated in a bi-parental cross of the cultivated octoploid strawberry (*F*. x *ananassa*). The mapping of resistance in controlled glasshouse experiments and the identification of QTL associated with resistance is reported. Furthermore, using historic crown rot disease score data, a genome-wide association study was conducted to investigate the presence of QTL within the wider germplasm. Subsequent to the identification of resistance QTL, the diploid strawberry reference genome (*F. vesca* Hawaii 4 v1.1) was mined for candidate resistance genes.

## MATERIALS AND METHODS

### Strawberry plant material and *Phytophthora cactorum* isolates

The mapping population used in this study was a cross between the cultivated June bearing strawberry cultivars ‘Emily’ x ‘Fenella’. ‘Emily’ is an early season variety with resistance to powdery mildew (*Podosphaera aphanis*), bred by NIAB EMR (formally HRI-East Malling) and released in 1995. It is moderately susceptible to *P. cactorum*. ‘Fenella’ is a mid-late season variety with good resistance to Verticillium wilt (*Verticillium dahliae*) and crown rot (*P. cactorum*), bred by NIAB EMR (formally East Malling Research) and released in 2009. The F_1_ full sib family of 181 individuals were clonally propagated by pinning down runners; the 181 progeny were planted in a field at the East Malling site in May 2014 and grown under netting. The strawberry runners were pinned down in beds and grown on for five months, from July 2014 – January 2015. The clones were then dug up, the excess soil was shaken off and the bare-rooted plants were transferred into a 2 °C cold-store for 1 week, before being transferred to a −2 °C cold-store for at least two months. Plants were brought out of cold-storage and potted into 9 cm diameter pots (Soparco) and dead leaves were removed. Plants were grown in a glasshouse compartment maintained at 20 ºC during the day and 15 ºC at night on a 16/8 hour, day/night light cycle, for three weeks before inoculation with *P. cactorum* isolates.

The main *P. cactorum* isolate used in this study for the bi-parental QTL mapping was P414. Isolates P404, P415 and P416 were used to screen the 15 representative (five ‘resistant’, five ‘intermediate’ and five ‘susceptible’) genotypes from the bi-parental cross. The isolates used in the genome-wide association study were P371, P372, P404, P407, P412, P413 and P416. All isolates are known to be pathogenic to *F.* x *ananassa,* having been isolated from infected strawberry plants. Isolates of *P. cactorum* were maintained on V8-juice (Arnotts Biscuits Limited) agar (200 ml V8-juice, 8-9 ml 1M KoH (Sigma-Aldrich), 20 g Agar (Fisher BioReagents) and 800 mL distilled water, autoclaved) at 20 ºC in the dark.

### *Phytophthora cactorum* zoospore production

Ten mm discs were cut from the margins of actively growing cultures of *P. cactorum* on V8-juice agar and placed into empty 9 cm triple-vented petri dishes (five per plate; Thermo Scientific). The plates were then carefully flooded with diluted compost extract (50 g compost in 2 L dH_2_O; and dH_2_O 1:1) and sealed with Parafilm (Bemis Company). Plates were placed in an incubator set at 20 ºC with lights on continuously for 48 hours to stimulate sporangia development. After 48 hours, the diluted compost extract was poured off and replaced with fresh diluted compost extract. The plates were placed into a fridge (~4 ºC) for 45 min and then moved onto the bench at room temperature for 45 min. The inoculum suspension was then vacuum filtered and kept on ice. The concentration of zoospores was calculated using a haemocytometer and the concentration was adjusted to 1 × 10^4^ zoospores per mL ^34^.

### Strawberry inoculation assays

The pathogenicity screens were carried out under controlled conditions in a glasshouse. Compartments were maintained at 20 ºC during the day and 15 ºC at night on a 16/8 hour, day/night light cycle for four weeks after inoculation with *P. cactorum*. The ‘Emily x ‘Fenella’ progeny screens were performed in six experiments. The first two experiments comprised of one replicate mock inoculated and one replicate artificially inoculated with *P. cactorum* isolate P414. The other four experiments were comprised of two replicates artificially inoculated with *P. cactorum* isolate P414. The 15 representative (five ‘resistant’, five ‘intermediate’ and five ‘susceptible’) genotypes screened with three other *P. cactorum* isolates (P404, P415 and P416) were carried out in a separate experiment and inoculated separately. Plants for all screens were arranged in a randomised block design for each set of replicates. In total, ten replicates of each strawberry genotype were artificially inoculated with a suspension of *P. cactorum* zoospores. Fifteen mm wounds were made using a scalpel on one petiole per plant and strawberry plants were sprayed with ~5 mL of 1 × 10^4^ zoospore suspension. For each strawberry genotype, two plants were mock inoculated by wounding in the same way and inoculated with ~5 mL diluted compost extract. To maintain humidity, plants were completely covered with clear plastic sheeting for 48 hours. Plants were scored following a slightly modified version of Bell *et al.*’s disease scale ^35^. Foliage was assessed visually for the presence of wilting symptoms once a week. The scores 8, 7, 6, 5 were assigned if the plant died during the first, second, third or fourth week after inoculation, respectively. After four weeks, the plants were cut open longitudinally and the crowns were assessed on a scale of 1-5; 1 – healthy (0% infection), 2 – up to 25% infection, 3 – 26-50% infection, 4 – 51-75% infection, 5 – 76-100% infection.

### Analysis of disease scores

The data for the ten replicates of each genotype was averaged and a mean crown rot disease score was used for further analysis. Statistical analyses were performed using R (v3.2.2, “Fire Safety” ^36^). The mean crown rot disease data for the ‘Emily’ x ‘Fenella’ progeny was tested for normality using the Shapiro-Wilk normality test. Broad sense heritability (*H*^*2*^) was calculated, *H*^*2*^ = *V*_*G*_/*V*_*P*_, where V_G_ is the total genetic variance and *V*_*P*_ is the total phenotypic variance.

### DNA extraction and genotyping

Young emerging leaf samples were collected in 2 mL microcentrifuge tubes along with two ball bearings and flash frozen in liquid nitrogen. Frozen leaf samples were ground to a fine powder for 2 mins at 60 o/m using a TissueLyser (Qiagen). Genomic DNA (gDNA) was extracted using the DNeasy kit (Qiagen) following the manufacturer’s protocol and eluted in 60 μL Buffer AE. gDNA quantity and purity were determined using the NanoDrop (ND-1000, Thermo Scientific) spectrophotometer. gDNA of ‘Emily’ and ‘Fenella’, the 181 progeny and 59 strawberry accessions, were sent to Oxford Genomics Centre for genotyping on the Affymetrix® IStraw90 Axiom® Array ^24^. Later, a further 55 strawberry accessions were genotyped on the Affymetrix® IStraw35 Axiom® Array ^26^.

### Linkage analysis of the bi-parental cross of ‘Emily’ x ‘Fenella’

Initial genotype calls were made using Affymetrix Power Tools (version 1.16.1) and the R package SNPolisher (version 1.5.0). Further filtering used custom Python scripts, part of the Crosslink package (https://github.com/eastmallingresearch/crosslink) to remove markers with strong segregation distortion ^37^. A bi-parental genetic map of ‘Emily’ x ‘Fenella’ was produced using the 181 progeny using the Crosslink software ^37^. The same pipeline was also used to generate bi-parental maps from IStraw90 data from four additional crosses: ‘Redgauntlet’ x ‘Hapil’, ‘Flamenco’ x ‘Chandler’, ‘Capitola’ x ‘CF1116’ (INRA) and ‘Camarosa’ x ‘Dover’ (CRAG). Custom Python and R scripts were used to create a consensus genetic map from all five bi-parental maps. Further custom scripts adjusted the fine scale marker ordering of the consensus map to match the *F. vesca* genome v2.0 ^22^ whilst identifying and correcting probable *F. vesca* genome assembly errors. The resulting hybrid consensus map was used to inform the ordering of the ‘Emily’ x ‘Fenella’ map.

The four sub-genomes of *F*. x *ananassa* were assigned the letters A, B, C and D in the ‘Emily’ x ‘Fenella’ linkage map. The letter denotes the similarity of the sub-genome to *F*. *vesca*, as described by van Dijk *et al.* ^38^. The most similar sub-genome was named A, the second most similar was named B (similar to the wild diploid *F. iinumae*), the third most similar was named C and the least similar was named D.

### QTL mapping

Histograms of mean crown rot disease scores were visualised and were tested for normality (QQ-plot). The raw mean data (*W*=0.96877, *p*<0.0004) was used for QTL analysis. QTL mapping was performed using Kruskal-Wallis (K-W) non-parametric ANOVA on the combined map of ‘Emily’ and ‘Fenella’. Identification of QTL specific to one parent and QTL that are present in both parental genotypes were estimated with the K-W ANOVA method, eliminating the need to perform separate QTL analysis on the two parental linkage maps. K-W analysis identifies markers linked to single traits/QTL individually and produces a *K* statistic. QTL associated with resistance to *P. cactorum* were identified if *p*<0.01, and the most significant marker was selected. A permutation test consisting of 10 000 iterations of randomly selected data was performed to verify that significance selection was not needed.

A stepwise linear regression model was performed to estimate the effect of each QTL. Non-significant QTL were removed from the model (*p*>0.05). The combination of the QTL effect sizes were used to estimate predicted means for each individual. The predicted means were plotted against observed average scores and their coefficient of determination (*r^2^*) was calculated. One-way analysis of variance (ANOVA) was performed to test for epistatic interactions between the three major effect QTL.

The Bonferroni correction was applied to the K-W *p* values to control the false-positive (type I error) rate. It was calculated using: critical *p* value (α)/number of comparisons being made. Only QTL with *p* values more significant than the Bonferroni correction were investigated further.

### Genome-wide association study

A preliminary genome-wide association study was performed using historic crown rot score data of strawberry accessions, collected between 1995-2017, using a mixture of two *P. cactorum* isolates each year: pre-2006 - isolates P371 and P372, 2006-2009 - isolates P412 and P413, 2010-2011 - isolates P412 and P407 and 2012-2017- isolates P404 and P416.

SNPs showing at least 5% minor allele frequency were assessed for association with resistance to crown rot in the 114 strawberry accessions using both PLINK ^39^ and TASSEL ^40^ (*p*<0.00005; https://github.com/harrisonlab/popgen/blob/master/snp/gwas_quantitative_pipeline.md). Population structure was taken into account and the Benjamini-Hochberg Procedure was used to reduce the false discovery rate and *p* values that lower than *p*=0.05 were considered to be potential QTL.

### Mining of candidate resistance genes in *Fragaria vesca*

The most significant SNP markers for each QTL were plotted on the *F. vesca* Hawaii 4 v1.1 genome ^41^ in Geneious (v10.1.2), along with GFF (generic feature format) files with the positions of NLR, RLK and RLP gene models. The number of genes in each of these classes within 1 Mbp either side of the most significant marker for each QTL were determined.

## RESULTS

### Variation observed in resistance to *Phytophthora cactorum* in the bi-parental cross

Crown rot disease severity was found to vary in a genotype-dependent manner. In the most susceptible individuals, total plant collapse occurred one or two weeks after inoculation and 100% necrosis of the crown was observed (crown rot disease scores 7/8, respectively). The mock-inoculated plants remained disease-free. The distribution of crown rot disease severity for the 181 individuals of the ‘Emily’ and ‘Fenella’ mapping population had a unimodal distribution pattern, with a slight skew towards resistance (Figure. S1). The means of the raw data were normally distributed and were used for QTL mapping of loci involved in resistance to *P. cactorum* (Figure. S1). The broad sense heritability was calculated to be *H*^*2*^ = 0.58.

### Linkage mapping

A whole-genome linkage map comprising of 11 598 SNP markers was assembled using the IStraw90 markers of the ‘Emily’ x ‘Fenella’ progeny and the program Crosslink ^37^. The map was resolved into 28 linkage groups, representing the four sub-genomes of each, of the seven chromosomes of *F.* x *ananassa*. Using only the IStraw35 markers to produce the map, 8 348 SNP markers were assembled into 28 linkage groups.

### QTL analysis

QTL mapping revealed 15 QTL significantly associated with resistance to *P. cactorum* isolate P414, located on chromosomes 1, 2, 3, 5, 6 and 7 (*p*<0.01; Figure 1a and Table 1). The most significant marker associated with each QTL is shown in Table 1. Comparing the K-W analysis between the markers from the IStraw90 array and the subset on the IStraw35 array, the same 15 QTL are still significant (*p*<0.01; Figure 1b and Table 1) before stepwise linear regression was performed. Only one focal SNP changed; IStraw35 SNP NE5FF79A1403E4 (2 868 559 bp) became the most significant marker on LG1B as the IStraw90 marker N5192084E36892 (2 433 670 bp) was removed, a distance of 434 889 bases is between the two markers on the *F. vesca* Hawaii 4 v1.1 genome ^41^ (Table 1).

**Figure 1.**
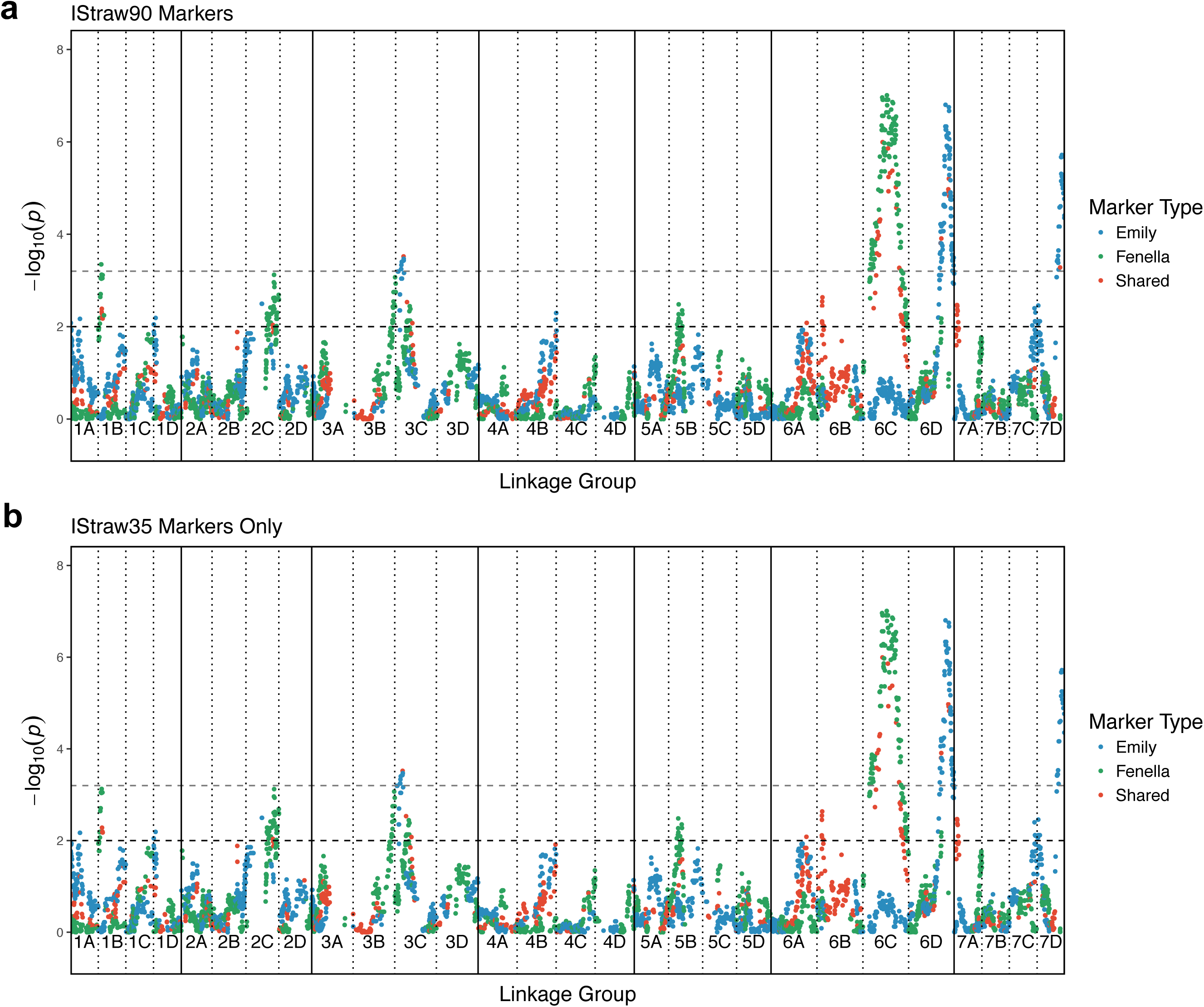
Quantitative trait loci analysis (QTL) results for ‘Emily’, ‘Fenella’ and shared markers across the 28 strawberry linkage groups. The coloured dots represent -log_10_(*p*) scores for the QTL analysis, while the black dashed horizontal line represents the significance threshold 2 (*p*=0.01) and the grey dashed horizontal line represents the significance threshold 3.2 (*p*=0.0006; Bonferroni correction). Regions on linkage groups 1A, 1B, 1D, 2C, 3B, 3C, 5B, 6A, 6B, 6C, 6D, 7A, 7C and 7D were significantly associated with resistance to *Phytophthora cactorum*, based on *p*<0.01. (**a**) IStraw90 markers, (**b**) IStraw35 markers only.

**Table 1.**
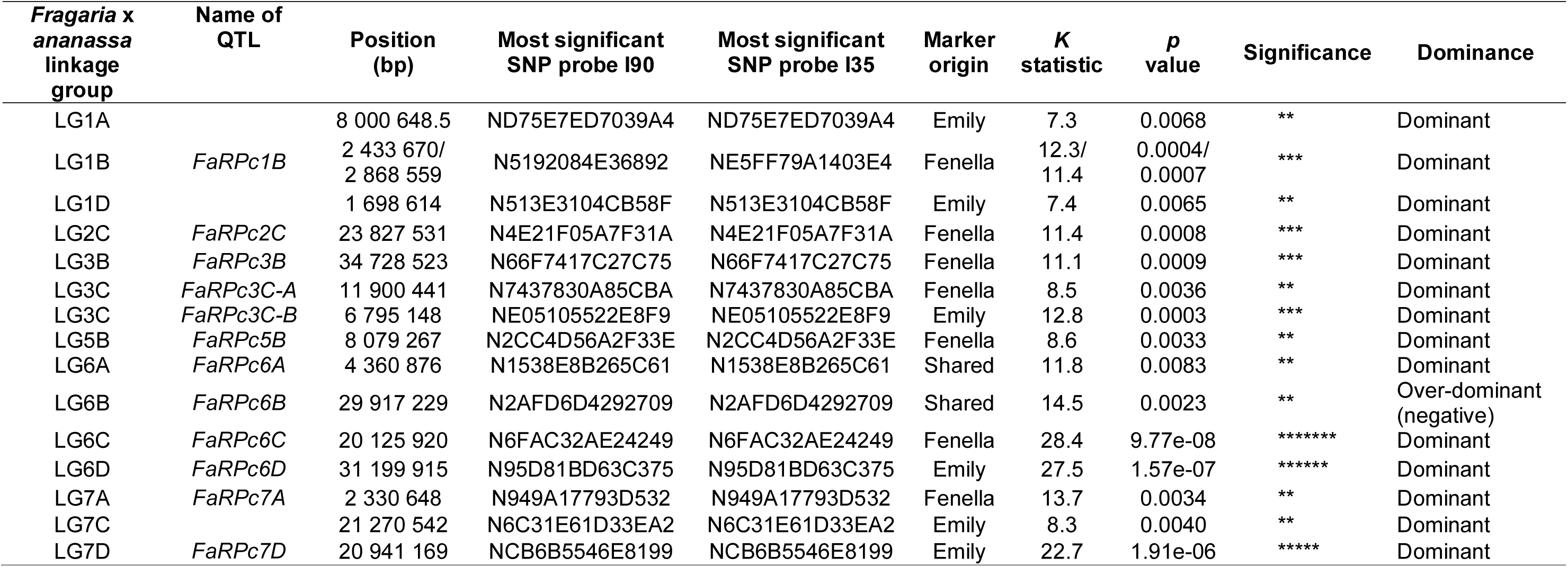
*Phytophthora cactorum* resistance quantitative trait loci (QTL) identified in the cultivated strawberry ‘Emily’ x ‘Fenella’ progeny by Kruskal-Wallis analysis using the Axiom^®^ IStraw90 and IStraw35 SNP arrays, *p*<0.01.

Stepwise linear regression was performed and three of these QTL were found to be non-significant, leaving 12 QTL associated with resistance to *P. cactorum* (Table S1). Seven QTLs were present in ‘Fenella’ only; named *FaRPc1B* (*<u>F</u>ragaria* x *<u>a</u>nanassa* <u>R</u>esistance to *<u>P</u>. <u>c</u>actorum* linkage group <u>1B</u>), *FaRPc2C*, *FaRPc3B*, *FaRPc3C-A*, *FaRPc5B*, *FaRPc6C* and *FaRPc7A*, located on linkage groups 1B, 2C, 3B, 3C, 5B, 6C and 7A, respectively (Table 1). Three QTLs were present in ‘Emily’ only; named *FaRPc3C-B*, *FaRPc6D* and *FaRPc7D*, located on linkage groups 3C, 6D and 7D, respectively (Table 1). Two QTL were present in both ‘Emily’ and ‘Fenella’, named *FaRPc6A* and *FaRPc6B*, located on linkage groups 6A and 6B, respectively (Table1). Of these significant resistance QTL, two were identified on the A sub-genome (the most similar to *F. vesca*), four resistance QTLs were identified on the B sub-genome (the most similar *F. iinumae*), four resistance QTLs were identified on the C sub-genome and two resistance QTLs were identified on the D sub-genome.

All but one of the QTL behave in a dominant nature. *FaRPc6B* is over-dominant in nature (Table 1). If the individual is homozygous for either parent (AA or BB), it has a predicted crown rot disease score of 2.8/2.7. However, if the individual is heterozygous (AB/BA) at that locus then it has a predicted crown rot disease score of 3.4/3.2. The individuals that are heterozygous at this locus are more susceptible than either homozygous individuals.

The linear regression also allowed the calculation of a contributing crown rot disease score to each of the 12 significant resistance QTL (Table S1). The estimate of effect sizes were combined to produce each individual’s predicted score (Table S2). Individuals with no resistance QTL were predicted to have a crown rot score of 5.237 (Table S1). The detected QTL explained 85.7% of the genetic variation observed; the combination of all remaining 12 QTL explain a reduction in crown rot score of 4.527, leaving a score of 0.71 unexplained by genotype within this analysis. The percentage effect of the 12 individual QTLs ranged from 4.2 – 13.3% (Table S1). The three largest effect QTL, *FaRPc6C*, *FaRPc6D* and *FaRPc7D*, accounted for 10.1%, 13.3% and 12.4%, respectively and explained a total of 35.8% of the variation observed. The predicted means were plotted against observed average scores and were found to be positively correlated, *r^2^* = 0.63 (Figure 2).

**Figure 2.**
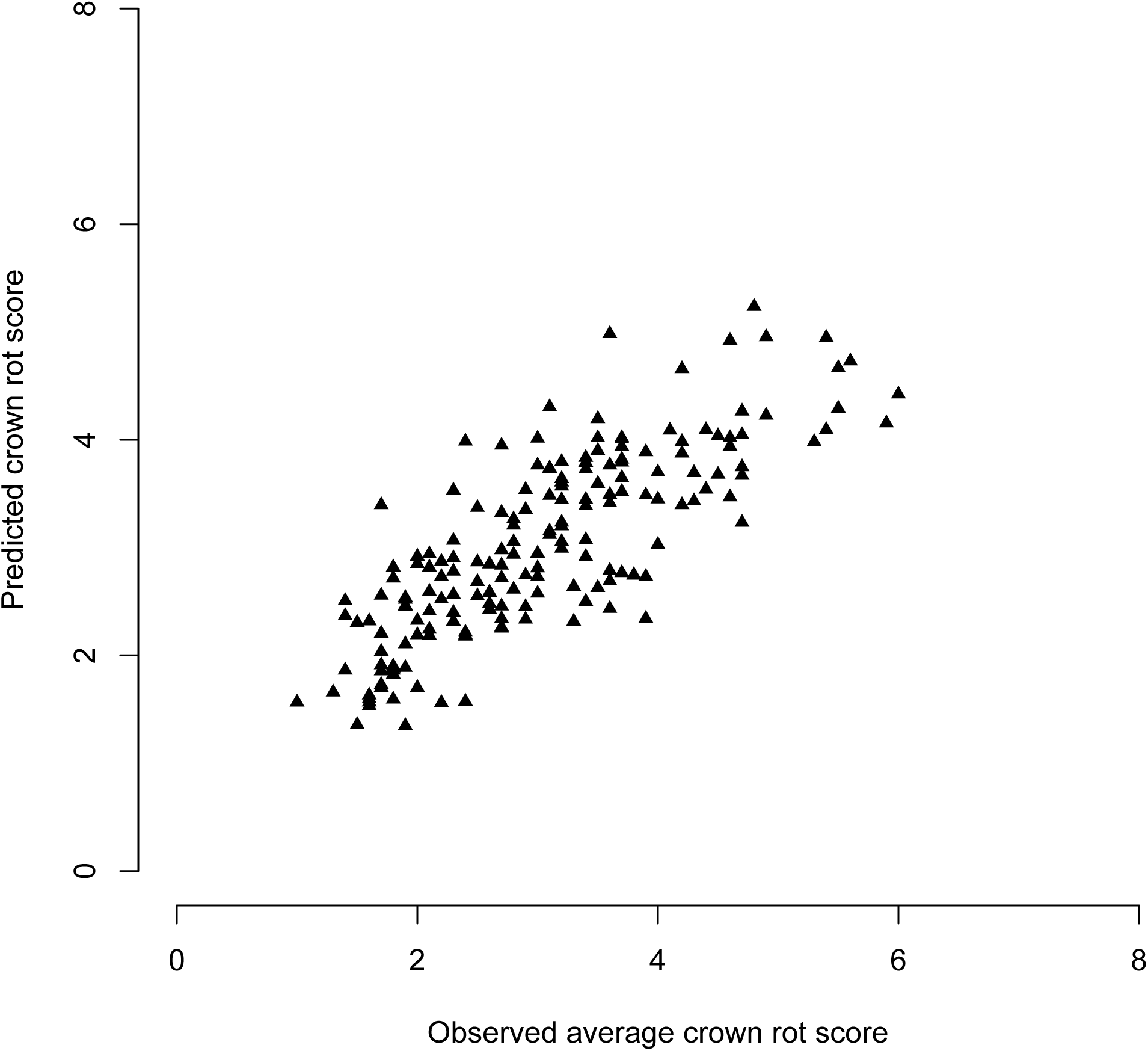
Predicted means are highly correlated with observed average crown rot scores, *r^2^* = 0.63.

The Bonferroni correction was calculated to be 0.01/15 = 0.00066 and plotted as - log_10_(0.00066) on the QTL analysis results (Figure 1a and 1b, grey dashed lines). Using the IStraw90 markers, five QTL cross this threshold; *FaRPc1B*, *FaRPc3C-B*, *FaRPc6C*, *FaRPc6D* and *FaRPc7D*, whereas only four cross the threshold using only the IStraw35 markers; *FaRPc3C-B*, *FaRPc6C*, *FaRPc6D* and *FaRPc7D* (Figure 1a and 1b). Focusing on the three major effect QTL, one-way ANOVA revealed there are no epistatic interactions between these QTL, *FaRPc6C*, *FaRPc6D* and *FaRPc7D*. Each marker has an effect on its own, is additive in nature and there are no pairwise interactions (Table S3).

### Analysis of the representatives of the ‘Emily’ and ‘Fenella’ population and their response to three further *Phytophthora cactorum* isolates

A subset of 15 individuals from the progeny; five resistant individuals (EF011, EF021, EF101, EF147 and EF184) with average disease scores ranging from 1 to 2, five intermediate response individuals (EF041, EF060, EF141, EF164 and EF187), with average disease scores ranging from 2.7 to 4.3 and five susceptible individuals (EF035, EF040, EF084, EF120 and EF166) with average disease scores ranging from 5.2 to 5.8, were tested for their response to three further isolates of *P. cactorum*, P404, P415 and P416 (Table 2). No major differences in host response were observed, individuals possessing few QTL were more susceptible than individuals possessing multiple QTL. However, some lines showed higher susceptibility than predicted (Table 2). Genotypes that possessed the three major effect QTL (*FaRPc6C*, *FaRPc6D* and *FaRPc7D*) or combinations of them were more resistant than those genotypes that did not possess any (Table 2).

### Presence of known classes of resistance genes in and around QTL regions and genome-wide association study

Focusing on the three major effect QTL (*FaRPc6C*, *FaRPc6D* and *FaRPc7D*) identified from the bi-parental cross, at least one NLR gene was identified within 1 Mbp either side of the most significant SNP, located on *F. vesca* Hawaii 4 v1.1 genome ^41^ (Table 3). At least one RLK was also identified in the same regions, as well as six RLP genes in the *FaRPc6D* region (Table 3).

**Table 3.**
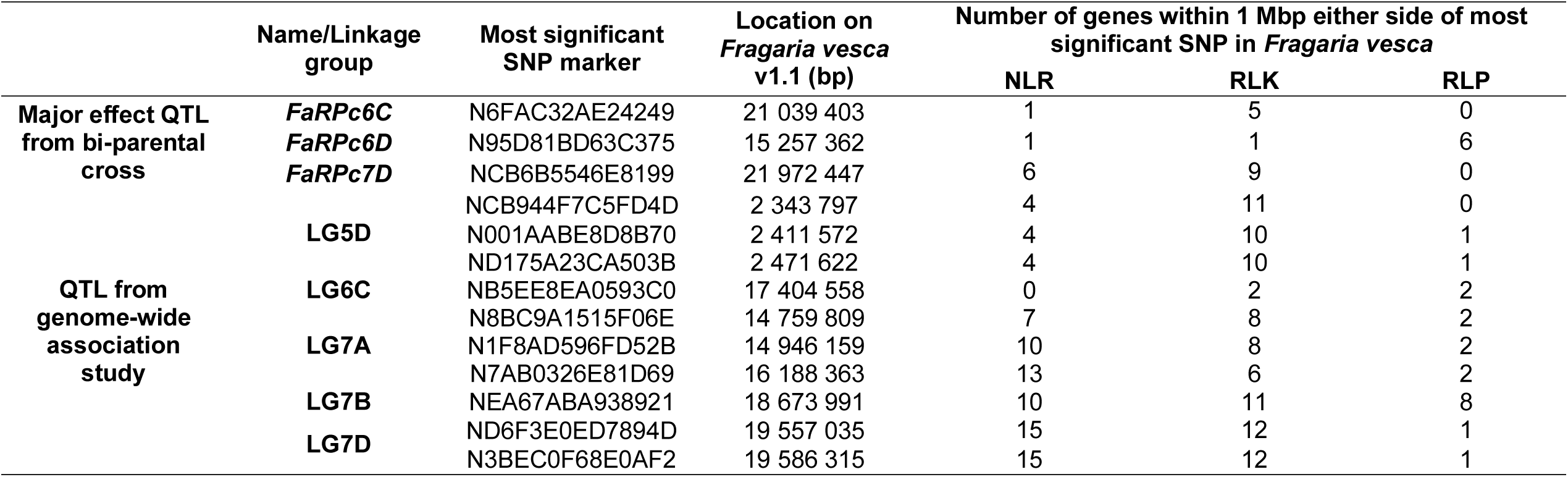
Details of the number of nucleotide-leucine rich repeat (NLR) receptors, receptor-like kinases (RLK) and receptor-like proteins (RLP) within 1 Mbp either side of the most significant marker on *Fragaria vesca* Hawaii 4 v1.1.

A genome-wide association study identified 10 SNPs significantly associated with resistance to *P. cactorum* on linkage groups 5D, 6C, 7A, 7B and 7D (Table 4). None of these markers overlapped with the QTL discovered in the bi-parental cross, although all linkage groups except 7B had QTL on them. The two SNPs identified on LG7D (ND6F3E0ED7894D and N3BEC0F68E0AF2) are located within the QTL region of *FaRPc2* identified by Mangandi *et al.* ^19^, based the on the positions on the *F. vesca* Hawaii 4 v1.1 genome ^41^. There is ~0.1 Mbp between the three SNPs identified on LG5D, with two RLKs located in this region. There is an RLP close to the SNP identified on LG6C (NB5EE8EA0593C0) and this SNP is ~7 Mbp upstream from *FaRPc6C* identified in the bi-parental cross. There were three SNPs identified on LG7A, two were close together and the was a RLK gene between them (N8BC9A1515F06E and N1F8AD596FD52B). There is also an RLK gene close to the other marker (N7AB0326E81D69).

**Table 4.**
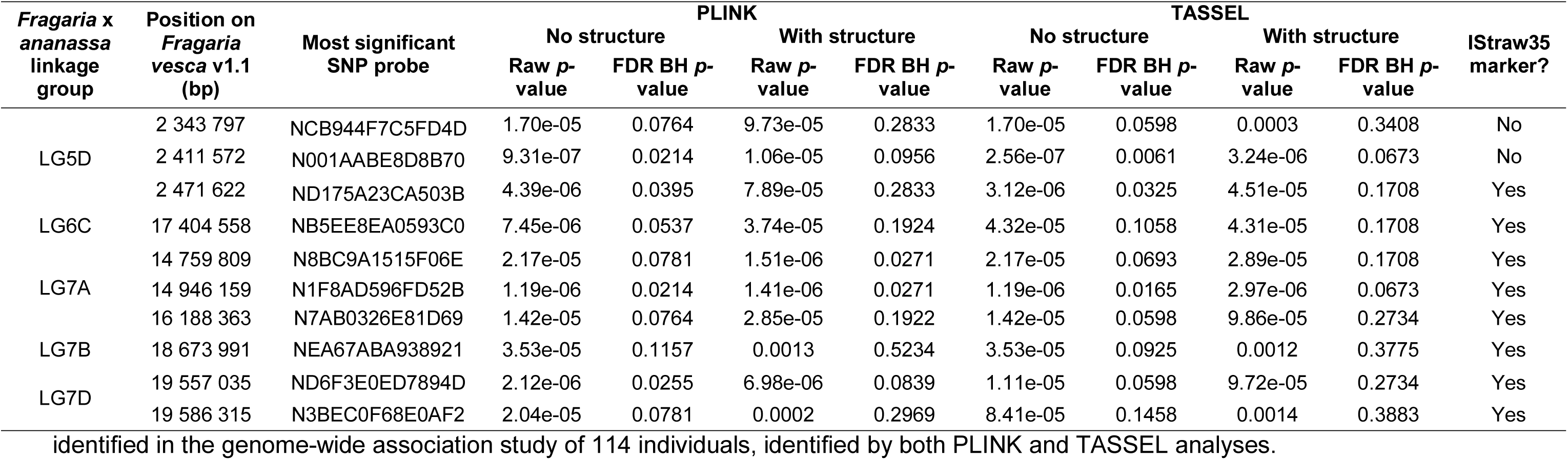
Details of most significant single nucleotide polymorphism (SNP) markers associated with resistance to *Phytophthora cactorum*

## DISCUSSION

Resistance to *P. cactorum* in octoploid strawberry is known to be under complex genetic control, with multiple QTL involved in resistance ^16,18^. In this study, single QTL provide small effects, but combined, in the bi-parental study explain 86% of the variation observed. A total of 15 putative QTLs for resistance to crown rot were identified in this study, of which five were found to still be significant after Bonferroni correction (*FaRPc1B*, *FaRPc3C-B*, *FaRPc6C*, *FaRPc6D* and *FaRPc7D*). In a previous study, five putative QTLs were identified in the *F. x ananassa* ‘Capitola’ x ‘CF1116’ progeny, with the QTL effects ranging from 6.5 to 10.2% and coming from both parents ^16^. Our results are comparable, with the QTL identified from both parents and with effect sizes ranging from 4.2 to 13.3%. The three major effect QTL, *FaRPc6C*, *FaRPc6D* and *FaRPc7D*, had an effect size of 10.1%, 13.3% and 12.4%, respectively and were highly significant with *p* values of 9.77e-08, 1.57-e07 and 1.91-e06, respectively.

Comparing the QTL analysis of the bi-parental cross using markers from the IStraw90 and IStraw35 arrays, there were no differences between the QTL identified, just a difference between the focal SNP for LG1B. The informative markers from this cross were used in the development of the IStraw35 array, along with informative markers from many other bi-parental crosses ^26^. The cheaper IStraw35 array provides sufficient markers to identify resistance QTL within the bi-parental cross as well as in the genome-wide association study, as most of the markers were present on the IStraw35 array.

Resistance (*R*) genes effective against *Phytophthora* typically contain NLR domains, which can directly or indirectly perceive pathogen effectors ^42^. NLR proteins are abundant in most plant genomes and are involved in the detection of a wide range of pathogens, oomycetes, fungi, viruses, bacteria, nematodes and insects. The diploid *F. vesca* has been well studied and at least 144 NLR genes have been identified ^43^. Along with this class of resistance gene, RLK and RLP genes have been associated with resistance to *Phytophthora* pathogens ^29–33^. Within 1 Mbp either side of the most significant marker of the three major effect QTL (*FaRpc6C*, *FaRPc6D* and *FaRpc7D*) from the bi-parental cross there is at least one NLR candidate gene in the *F. vesca* genome. There is also at least one RLK candidate gene within 1 Mbp *FaRpc1B*, *FaRpc3C-B*, *FaRpc6D*, and *FaRpc7D* and multiple RLP candidate genes within and around *FaRpc1B*, *FaRpc3C-B*, *FaRpc6D*, and *FaRpc7D*. Disease resistance genes are often found in clusters in the genome ^44^. Further work is required to identify the genes underlying the resistance QTL.

No race structure has been reported for *P. cactorum*. The consistency between the response of the 15 progeny to the four isolates of *P. cactorum* is promising in that the resistance identified in the ‘Emily’ x ‘Fenella’ population would be useful against different isolates of *P. cactorum*. If the three major effect QTL (*FaRpc6C*, *FaRpc6D* and *FaRpc7D*; combined score reduction of 2.016) could be introgressed into elite strawberry germplasm, then a good base level of resistance to *P. cactorum* could be provided, with the other QTL, such as *FaRpc1B* supplementing them. Interestingly ‘Emily’ possesses two of the major effect QTL (*FaRPc6D* and *FaRPc7D*), but is more susceptible than ‘Fenella’. Possibly there are additional undetected factors affecting the overall resistance/susceptibility status of the plant. Similarly, some genotypes showed higher susceptibility than predicted (EF120 and EF167; Table 2) indicating that additional undetected factors may affect the expression of some QTL.

In *F. vesca* a single major gene locus was identified on the proximal end of linkage group 6, named *RPc-1* (*<u>R</u>esistance to <u>P</u>hytophthora <u>c</u>actorum <u>1</u>*). *RPc-1* explained 74.4 % of the variation observed and was identified as spanning a region of 801 genes and 69 potential plant disease resistance genes, including multiple different classes of resistance gene, in a 3.3 Mb region of linkage group 6 from position 5 151 532 to 9 201 791 bp ^45^. *RPc-1* would be on *F. x ananassa* LG6A, based on the convention of van Dijk *et al.* ^38^. In the bi-parental cross a minor QTL was identified on LG6A, imputing this on *F. vesca*, it is very far away, ~29 Mbp upstream of *RPc-1*. Toljamo *et al.* observed the expression of potential resistance genes in the *RPc-1* region, in *P. cactorum* inoculated *F. vesca* Hawaii 4 roots ^46^. Within the *RPc-1* locus, four NLR protein-encoding genes (101306457, 101297569, 101300750 and 101304699) were identified as being expressed, two of which were significantly down-regulated within the inoculated plants (101300750 and 101304699) ^46^. The authors propose 101297569 as a strong candidate as it had the highest expression (mean expression level, 32.41) of the NLR genes in that region. Other types of resistance genes were identified within the *RPc-1* locus, two L-type-lectin-RLKs (receptor-like kinase) were significantly upregulated (101310048 and 101309756; log_2_ fold change 3.50 and 2.25, respectively), these were both considered strong candidates. Two G-type-lectin-RLKs (101305393 and 101305094) and one RLP (101290881) were also upregulated in the inoculated roots ^46^. Further characterisation is required to fully understand the resistance mechanisms against *P. cactorum* in *F. vesca*, as well as *F*. x *ananassa*.

Two of our major effect QTL were identified on the D sub-genome (*FaRPc6D* and *FaRPc7D*), which is the least similar to *F. vesca* ^38^. Recently, a major locus associated with resistance to *P. cactorum* in *F.* x *ananassa* was identified on linkage group 7D, named *FaRPc2* ^19^. The small genome-wide association study of the wider germplasm highlighted different areas of the genome associated with resistance to *P. cactorum*, compared to the QTL identified from the bi-parental cross, indicating further loci, not captured in our study of ‘Emily’ and ‘Fenella’ to be exploited for resistance. Two of these SNPs were in the same region as the major locus *FaRPc2*, identified by Mangandi *et al.* ^19^. In the same study, further QTL on LGs 1D, 3B, 5B, 6A and 6B were also identified, however, they were not consistent across the replicates or significant enough to be confirmed ^19^.

Despite the genome-wide association study being underpowered due to the small number of individuals investigated, several significant SNPs were identified and overall there was good concordance between the two approaches used, PLINK and TASSEL, but some SNPs were more significant in one method, compared to the other. Further individuals are required to be tested to increase the power of the analysis.

Quantitative resistance provides many challenges to breeders due to the complex nature of inheritance. However, this complexity can increase the durability of resistance as it is harder for the pathogen to overcome ^47^. To a large extent, the ‘Emily’ x ‘Fenella’ progeny responded similarly to each of the isolates tested (though there is some slight indication of variation in some *P. cactorum* isolates to the same plant genotypes), indicating that the three major effect QTL would be useful in commercial cultivars to provide broad-spectrum partial resistance. In future studies, it would be useful to explore the resistance in the wider germplasm, using a larger genome-wide association study approach on many hundreds of accessions, using the same isolate. This would provide more information about the status of resistance within the population and identify parents with other desirable traits. Subsequent work is also required to identify the gene(s) within the QTL regions and elucidate the mechanism of resistance. Future work will address whether the quantitative resistance to *P. cactorum* is a combination of multiple weak gene-for-gene interactions between the host and the pathogen by studying both the host and the pathogen in greater detail.

## ACKNOWLEDGEMENTS

This research was supported by grants awarded to RJH from the Biotechnology and Biological Sciences Research Council (BBSRC - BB/K017071/1, BB/N006682/1). The authors acknowledge the East Malling Strawberry Breeding Club for access to strawberry material, Dr Beatrice Denoyes, INRA and Dr Amparo Monfort, CRAG for granting the use of their informative markers in the production of the strawberry consensus linkage map, as well as the Farm and Glasshouse staff of NIAB EMR for their assistance and support. The authors also acknowledge the help received from Mr Antonio Gomez-Cortecero, Dr. Helen M. Cockerton and Dr. Bethany P. J. Greenfield and the members of RJH’s group in the preparation and setting up of experiments.

## FIGURE LEGENDS

**Figure 3.**
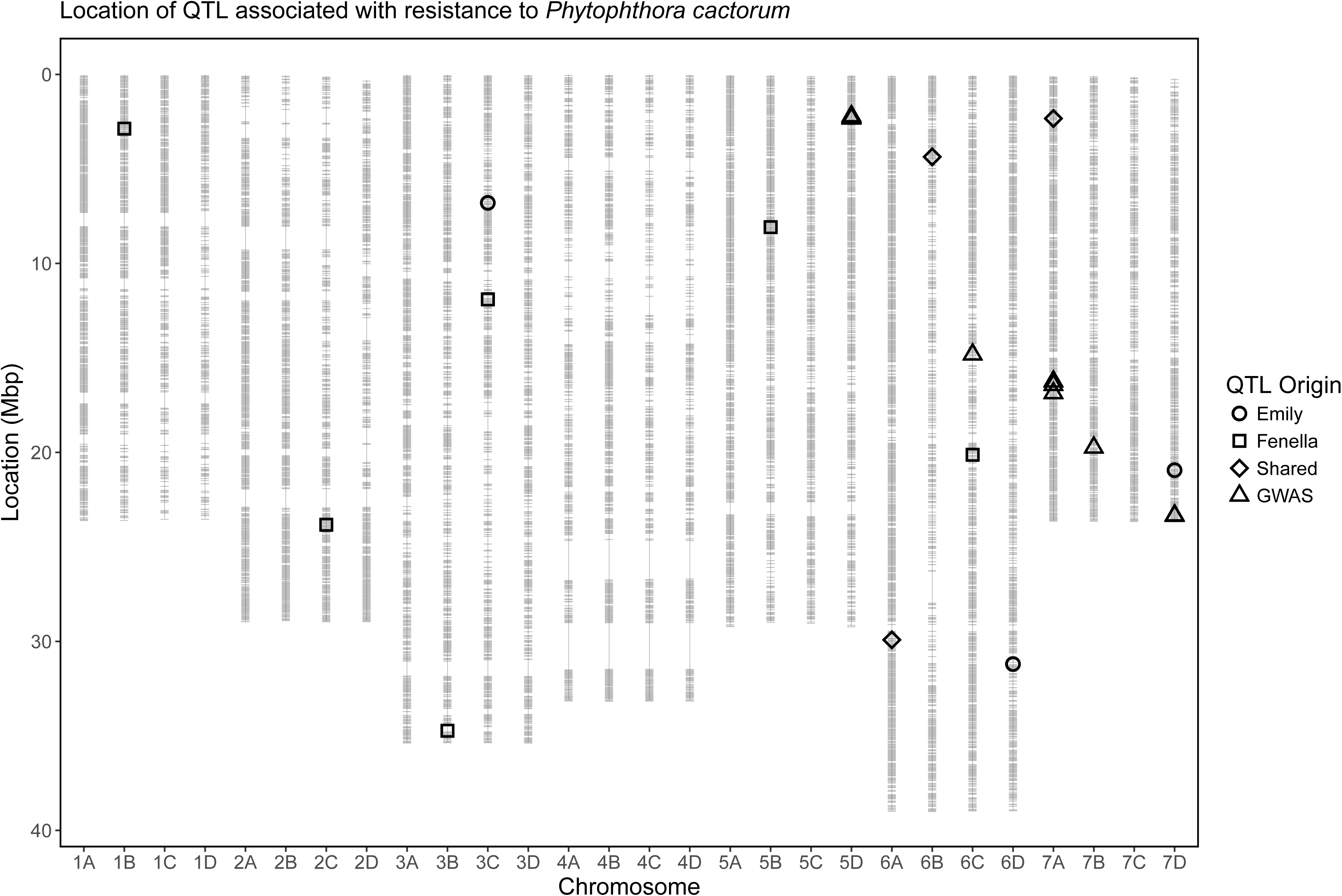
Combined map of ‘Emily’ x ‘Fenella’, depicting the position of *Phytophthora cactorum* resistance quantitative trait loci (QTL) from the bi-parental cross and the genome-wide association study. Circles are QTL from ‘Emily’, squares are QTL from ‘Fenella’, diamonds are QTL shared in both cultivars and triangles are QTL identified from the genome-wide association study (GWAS).

**Figure 4.**
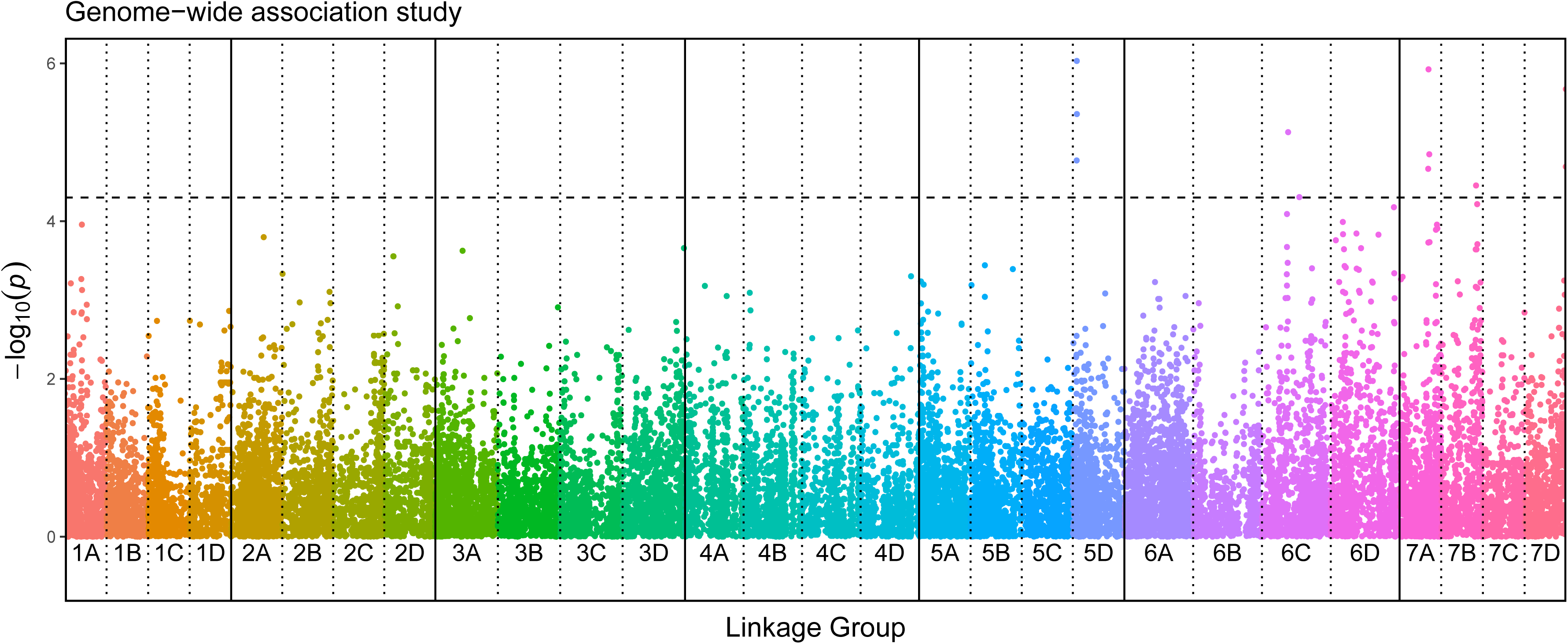
Manhattan plot of genome-wide association study on 114 strawberry accessions across the 28 strawberry linkage groups. The coloured dots represent - log_10_(*p*) scores for the QTL analysis, while the black dashed horizontal line represents the significance threshold 4.3 (*p*=0.00005). Regions on linkage groups 5D, 6C, 7A, 7B and 7D are significantly associated with resistance to *Phytophthora cactorum*.

## SUPPLEMENTARY FIGURE LEGENDS

**Figure S1.** Histogram of phenotypic distribution of average crown rot (*Phytophthora cactorum*) disease severity data for the ‘Emily’ x ‘Fenella’ progeny; ‘Fenella’ (2.1) and ‘Emily’ (4.4) average scores highlighted.

